# Predicting the Dynamic Viscosity of High-Concentration Antibody Solutions with a Chemically Specific Coarse-Grained Model

**DOI:** 10.1101/2025.09.17.676805

**Authors:** Tobias M. Prass, Patrick Garidel, Michaela Blech, Lars V. Schäfer

## Abstract

The viscosity of high-concentration protein solutions is a critical parameter in bio-pharmaceutical formulation development. Conventionally, the viscosity is measured and optimized in labor-intensive experimental workflows that require a lot of material. While predicting the viscosity with atomistic molecular dynamics (MD) simulations is feasible, they are computationally prohibitively expensive due to the large system sizes and the long simulation times involved. Coarse-grained MD (CG-MD) simulations significantly reduce computational demands, but evaluating their accuracy and predictive power requires rigorous validation. Here, we assess the capability of the Martini 3 CG force field to predict the viscosity of high-concentration antibody solutions. We show that a refined Martini 3 force field, with optimized protein–protein interactions, can predict the elevated viscosities observed in concentrated solutions of F(ab’)_2_ fragments of the therapeutic monoclonal antibody (mAb) omalizumab. Furthermore, we show that our previously developed Martini 3-exc model for arginine excipients successfully captures the trend of lowering viscosity, as observed in our rheology experiments. These findings open the way to physics-based computational prediction of the properties of dense biopharmaceutical solutions via large-scale MD simulations.

---

High-concentration protein solutions are central to numerous biological and pharmaceutical applications. In the context of biopharmaceuticals, high-concentration formulations are particularly desirable for subcutaneous injection, as they enable self-administration of therapeutic doses in small volumes. However, developing high-concentration formulations involves significant challenges, including issues related to protein conformational stability, particle formation, phase separation, and elevated viscosity;^1–6^ for a recent review, see Manning et al. ^7^ The crowded macromolecular environment in these solutions can influence biomolecular structure and solvation, and the kinetics of molecular processes.^8–13^

Monoclonal antibodies (mAbs), a major class of biopharmaceuticals, are especially affected by these factors due to their large size and conformational flexibility. mAbs are widely used in the treatment of diseases ranging from cancer and multiple sclerosis to asthma and COVID-19.The high therapeutic dosages required (up to 15 mg per kg body weight) and the limited volumes suitable for subcutaneous injection (up to about 3 ml)^15^ necessitate mAb concentrations of 100 mg/ml or higher. Elevated viscosity in these formulations is associated with increased injection forces, tissue back-pressure, and injection pain, while protein particles can decrease drug efficacy and trigger immunogenic responses. ^15,16^

Traditionally, strategies to optimize the stability and properties of biopharmaceutical solutions rely on empirical trial-and-error workflows that screen various combinations of buffers and excipients, such as salts and amino acids.^17,18^ Given the high production costs of mAbs,^19^ computational methods that can predict the viscosity of high-concentration formulations would be highly advantageous. Molecular dynamics (MD) simulations can, in principle, resolve the relevant molecular-level interactions underlying the properties of these systems. Our previous work demonstrated that all-atom MD simulations can correctly capture the increase in dynamic viscosity with rising mAb concentration. ^20^ However, the computational resources required for statistically robust results are enormous, limiting the practical application of such approaches.^20^ Alternative, more easily accessible methods include colloidal hard-sphere models^21–23^ or super-coarse grained (CG) representations that model mAbs with as few as, for example, 6, 12, or 24 CG beads and omit explicit solvent.^24–27^ Such computationally cheap models enable long time scale simulations of systems with a large number of mAb molecules. However, the reduced accuracy associated with the reduction of the proteins to a very coarse (sub-)domain level representation limits the predictive power of such super-CG models and necessitates careful parametrization and validation against experiments or higher-level simulations.

The molecular mechanisms governing the properties of high-concentration mAb solutions, as well as the viscosity-reducing and aggregation-mitigating effects of excipients, remain incompletely understood. Arginine, a widely used amino acid-based excipient, is known to mitigate viscosity and aggregation. ^28–30^ Recent atomistic MD studies have focused on on the effect of arginine on the Fab domains of mAbs, highlighting the interactions of arginine excipients with aromatic and charged amino acids.^31–33^ To directly investigate excipient effects on high-concentration solutions, a computationally efficient compromise between the super-CG and atomistic representation is desired – one that balances chemical specificity and sufficient sampling capability. Such a model should combine an explicit description of protein–excipient interactions with long-timescale simulations to sample all relevant configurations, in particular, the diverse protein–protein interaction interfaces and their modulation by excipient molecules.

The Martini CG force field is a widely used approach that achieves this balance by mapping fragments of two to four heavy atoms (plus the associated hydrogens) to a single CG bead. This level of resolution retains chemical specificity of protein–excipient interactions.^34,35^ The latest version, Martini 3, has improved interaction parameters compared to its predecessor Martini 2^36^ and can be used for large-scale multi-protein simulations. ^37–39^ However, the scale of the protein–protein interactions remains a potential shortcoming of Martini 3. For example, Thomasen et al. reported that Martini 3 yields too compact structures of intrinsically disordered protein (IDP) and multidomain proteins. They suggested to either increase the protein–water interactions or decrease the protein–protein interactions (PPIs) (by changing the well depths of the nonbonded Lennard-Jones (LJ) potentials by +10 % and -12 %, respectively).^40,41^ Binder et al. improved PPIs of lysozyme and subtilisin systems by applying separate scalings of nonbonded interactions between organic and ionic bead types.^42^ The Martini3-IDP force field addresses overly compact conformational ensembles of IDPs by adjusting the bonded interactions in the force field, improving agreement with experimental data.^43^ Pedersen et al. and Lamprakis et al. reported unstable protein complexes with Martini 3 that were stable in all-atom simulations.^44,45^ These findings suggest that the remaining inaccuracies in the model arise from factors that are more complex than a simple global over- or underestimation of protein–protein and/or protein–water interaction strength.

Dynamic (shear) viscosity can be computed from equilibrium MD simulations with a Green-Kubo (GK) formalism, which relates viscosity to the integral over the pressure autocorrelation function (ACF)^46,47^

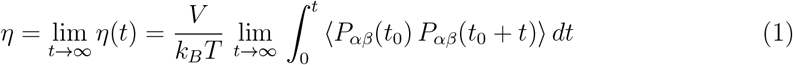

where *V* is the volume of the simulation box, *k*_*B*_ is Boltzmann’s constant, *T* is the temperature, and ⟨ *…*⟩ indicates thermal averaging. *P*_*αβ*_ are the nondiagonal elements of the pressure tensor, whose ACFs are calculated by averaging over all time origins *t*_0_. A major challenge of this method is sampling. The viscosity computed from the GK integral (Eq. 1) is exact in the asymptotic (long-time) limit, whereas the timescales accessible with simulations are limited (*µ*s up to maybe ms). Moreover, the large pressure fluctuations in the microscopic simulation systems result in slow convergence of the GK integrals. Large-scale all-atom MD simulations in explicit solvent have been used to study viscosity and diffusion in concentrated solutions of small proteins, which do not exhibit very high viscosities. ^48^ Our previous all-atom MD simulation study of a highly viscous mAb solution demonstrated that the GK approach can capture the experimental trend of viscosity changes with protein concentration.^20^ However, despite the huge computational resources required for achieving the multi-*µ*s time scales in the all-atom simulations, the statistical precision of the computed viscosity predictions was limited, and only a single system could be investigated.^20^ Furthermore, only small simulation systems containing four mAb molecules could be studied in our previous all-atom simulations.

In this work, we combine MD simulations and rheology experiments to assess the capability of the Martini 3 CG force field to predict the viscosity of high-concentration mAb solutions. Specifically, we address three key questions: (1) How does scaling of protein– protein interactions in Martini 3 affect viscosity predictions? (2) Can Martini 3 describe the increase in viscosity with rising mAb concentration? (3) Can the model capture the experimentally observed modulation of viscosity by arginine excipients? We use the therapeutic antibody omalizumab (Xolair^®^) as a model system, as its high-concentration solutions exhibit pronounced viscosity increases in the absence of excipients and substantial viscosity reduction upon addition of arginine chloride (Arg/Cl).

To facilitate MD simulations, we used F(ab’)_2_ fragments rather than full-length mAbs (Figure 1). This approach circumvents the need to model the glycosylated Fc domain of the mAb, which is a challenge because carbohydrate parameters for glycosylated amino acids are still under development and are not yet available in Martini 3.^49–51^ Furthermore, the smaller system sizes enable longer simulation times and larger simulation systems. We simulated systems containing 32 omalizumab F(ab’)_2_ fragments to investigate the viscosity of the high-concentration solutions (Figure 1). We tested different PPI scalings in Martini 3, including the 12% scaling of the well-depths of the Lennard-Jones interactions between protein beads suggested by Thomasen et al. for IDPs,^40,41^ and compare the resulting viscosities to experimental values. We also incorporated our previously developed arginine excipient model for Martini 3^52^ to evaluate whether the refined CG model captures the viscosity difference between Na/Cl and Arg/Cl environments. Finally, we probed mAb concentrations of 150, 200, and 250 mg/ml to investigate concentration-dependent changes in viscosity. Corresponding rheology experiments were performed for omalizumab F(ab’)_2_ at different concentrations and in different excipient environments. Method details are provided in the Supporting Information.

**Figure 1.**
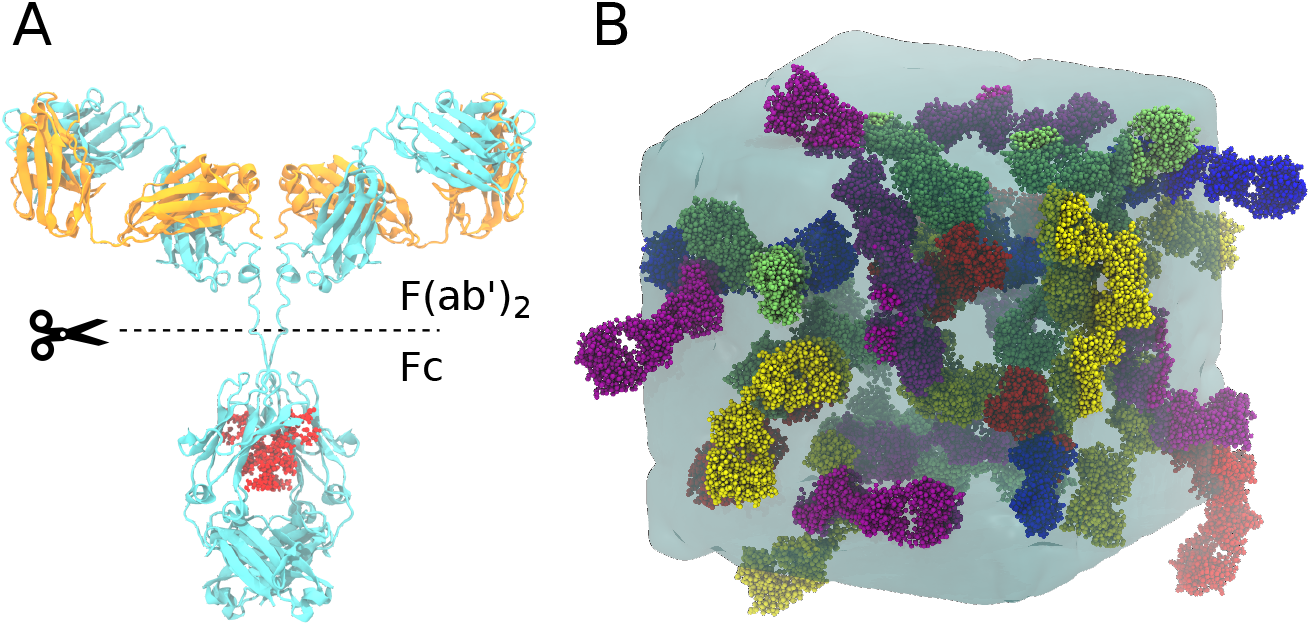
A) Molecular representation of full-length omalizumab. The cleavage into F(ab’)_2_ and Fc fragments is indicated by the dashed line. The heavy and light chains are shown in cyan and orange, respectively, and the glycan in the Fc domain is shown in red. B) Snapshot of one of the simulation systems containing 32 F(ab’)_2_ fragments at a concentration of 200 mg/ml.

We first analyzed simulations of F(ab’)_2_ systems at a concentration of 200 mg/ml (with 150 mM Na/Cl), using both the original Martini 3 force field parameters and a modified force field in which the protein–protein interactions were weakened by reducing the well-depths *ϵ*_*LJ*_ of the Lennard-Jones interaction potentials between protein beads. The scaling of *ϵ*_*LJ*_ was varied between 6% and 36%. For each system, five independent 5 *µ*s MD simulations were run (25 *µ*s in total), see Methods. The resulting Green-Kubo integrals and the final viscosity values are shown in Figure 2 and Table 1; GK integrals from the five individual repeat simulations are shown in Figure S1.

**Table 1:** Dynamic viscosities from the Martini 3 simulations of the 200 mg/ml F(ab’)_2_ systems with 150 mM Na/Cl and the different PPI scalings applied.

| PPI scaling | Viscosity (mPa · s) |
| --- | --- |
| 0% | 651.5 ± 301.7 |
| 6% | 196.9 ± 52.2 |
| 12% | 24.4 ± 6.5 |
| 24% | 5.4 ± 1.2 |
| 36% | 4.5 ± 0.4 |

**Figure 2.**
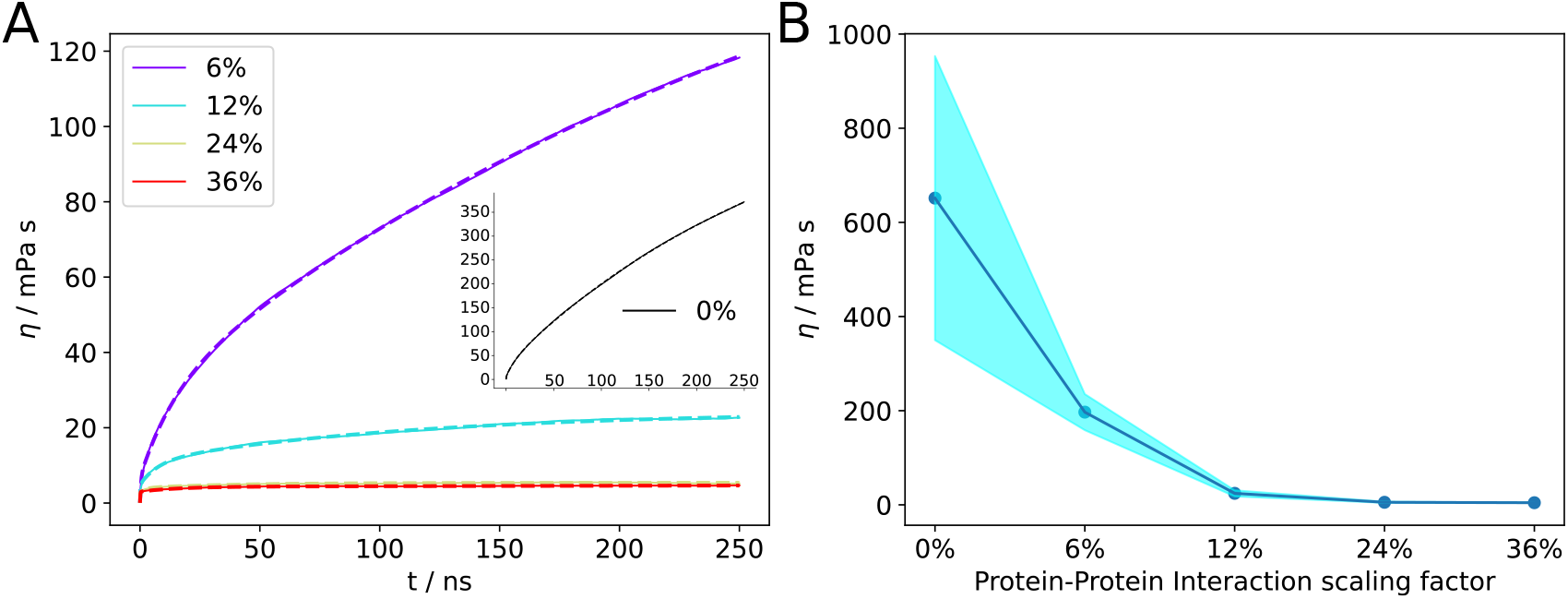
A) Green-Kubo integrals (solid lines) from the CG-MD simulations with the different PPI scalings. Exponential fits are shown as dashed lines. The inset in A) shows the results for the original (unscaled) Martini 3 force field. B) Dynamic viscosities obtained from the asymptotic limit of the exponential fits. The uncertainty (shaded area) represents the standard deviation of the computed values from the five individual repeat simulations.

The original Martini 3 force field yields a very high viscosity of 651 mPa · s, with a large statistical uncertainty due to the incomplete sampling of the ultraslow dynamics related to the reconfiguration of the sticky protein network in this highly viscous system. Weakening the attractive interactions between the protein CG beads by scaling the well-depths of the Lennard-Jones potentials strongly reduces the viscosity, with 6% and 12% PPI scalings yielding viscosity values of 196.9 and 24.4 mPa · s, respectively (Table 1). For comparison, the dynamic viscosity of pure Martini 3 water at 300 K is 0.66 mPa · s (Figure S2), which is about 20% lower than the experimental value of 0.853 mPa · s.

To experimentally validate the viscosity values predicted by the CG-MD simulations, we measured the viscosity of omalizumab F(ab’)_2_ fragments at different concentrations (Table 2). Our rheology experiments yielded viscosities of 12.6 and 37.1 mPa · s at the concentrations of 149 and 204 mg/ml, respectively. These values are in line with Heisler et al., who reported a viscosity of 19.3 mPa · s for a 180 mg/ml omalizumab F(ab’)_2_ solution.^53^

**Table 2:** Dynamic viscosity obtained from rheology experiments of different omalizumab F(ab’)_2_ concentrations in presence of 150 mM Na/Cl or 150 mM Arg/Cl (pH 6.5, 10 mM histidine buffer).

| Excipient conditions | F(ab')<sub>2</sub> concentration (mg/ml) | Viscosity (mPa · s) |
| --- | --- | --- |
| 150 mM Na/Cl | 13 | 1.9 ± 0.2 |
|  | 56 | 1.7 ± 0.4 |
|  | 103 | 3.4 ± 0.1 |
|  | 149 | 12.6 ± 0.3 |
|  | 204 | 37.1 ± 3.5 |
|  | 240 | 74.2 ± 2.1 |
| 150 mM Arg/Cl | 9 | 1.2 ± 0.3 |
|  | 48 | 1.3 ± 0.3 |
|  | 98 | 2.6 ± 0.1 |
|  | 152 | 7.9 ± 0.3 |
|  | 196 | 11.9 ± 1.2 |
|  | 245 | 26.7 ± 1.9 |

The discrepancy between the simulated and experimental viscosities suggests that improvements can be achieved by weakening the attractive protein–protein interactions in the Martini 3 force field. While it is possible to adjust the PPI scaling such as to quantitatively reproduce the experimental viscosity values, such an approach might be system-specific and would raise the risk of overfitting. Instead, we used the 12% scaling previously introduced by Thomasen et al. ^40,41^ for IDPs, accepting that the viscosities are slightly lower than experimental values (24.4 mPa · s at 200 mg/ml in the CG-MD simulation versus 37.1 mPa · s at 204 mg/ml in the rheology experiment). The primary goal of the CG-MD simulation approach is not necessarily to exactly reproduce the experimental numbers, but to predict trends, such as the effects of excipients (see below).

As a control, to investigate finite box size effects, we repeated the simulations at 200 mg/ml with a twofold larger system containing 64 F(ab’)_2_ fragments, yielding a dynamic viscosity of 20.0 ± 2.3 mPa · s (Figure S3) that is consistent with the value obtained with 32 F(ab’)_2_ fragments. Furthermore, the effect of pH was investigated by performing additional simulations at 12% PPI scaling in which all histidine side chains were doubly protonated (positively charged) to mimic low pH conditions in the pH range of approximately 5 to 6.5 (where the carboxylate groups of the acidic amino acids Asp and Glu are not yet protonated). The resulting viscosity is statistically indistinguishable from that obtained with standard histidine protonation states (Figure S4). As an additional control, to study the possible influence of the particular choice of the restraining potentials required to stabilize the secondary and tertiary structure of the Fab domains in the CG-MD simulations, we repeated the simulations of the 200 mg/ml F(ab’)_2_ systems with 150 mM Na/Cl with the recently published GōMartini 3 model^54^ instead of the Gō-like OLIVES potentials.^44^ Again, the viscosity value obtained from the simulations is statistically indistinguishable from the one with OLIVES restraints (Figure S4), further underlining the robustness of the findings.

We previously developed a new model for arginine excipients in Martini 3, which also involved reparametrizing the arginine–protein interactions at the single-residue level against reference data from all-atom simulations (coined the Martini3-exc model). ^52^ Since the excipient arginine is known to reduce the viscosity of omalizumab solutions,^55^ we used our arginine excipient model in conjunction with the F(ab’)_2_ systems to compare the viscosities between 150 mM Na/Cl and 150 mM Arg/Cl, thus testing the transferability and predictive capacity of the CG-MD approach. The viscosity data from the 150 mM Arg/Cl simulations are presented in Figure 3 and Table 3; the GK integrals of the five individual simulations are shown in Figure S5.

**Table 3:** Dynamic viscosities from the CG-MD simulations of the F(ab’)_2_ at 200 mg/ml in 150 mM Arg/Cl. Statistical uncertainties were estimated from the standard deviations of the five independent repeat simulations run for each system.

| PPI scaling | Viscosity (mPa · s) |
| --- | --- |
| 0% | 374.9 ± 151.9 |
| 6% | 20.0 ± 7.3 |
| 12% | 10.5 ± 1.7 |

**Figure 3.**
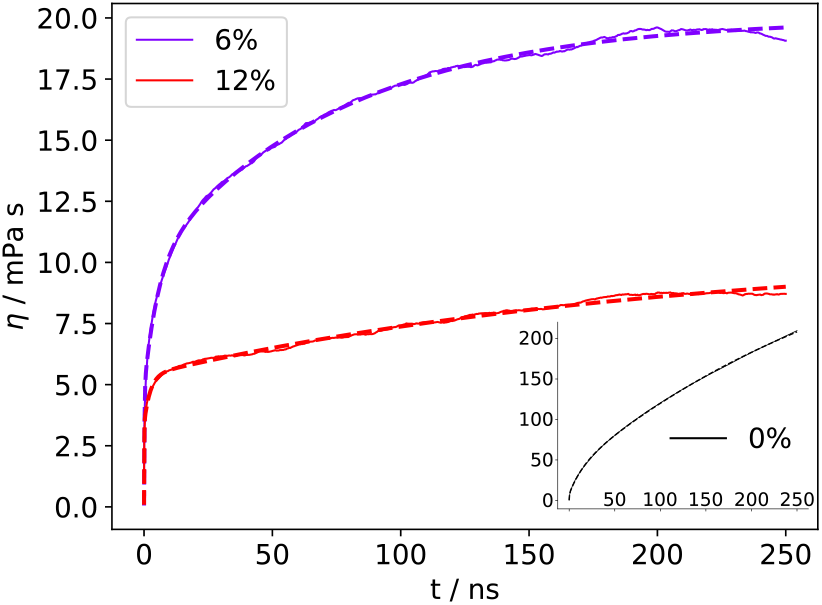
GK integrals (solid lines) and fits (dashed lines) from the CG-MD simulations of the F(ab’)_2_ at 200 mg/ml in 150 mM Arg/Cl.

In the presence of 150 mM Arg/Cl, the viscosity of the 200 mg/ml omalizumab F(ab’)_2_ system is reduced compared to 150 mM Na/Cl, by factors of approximately 1.7, 10, and 2.3 with 0%, 6%, and 12% PPI scaling, respectively (Tables 1 and 3). The observed viscosity reduction by a factor of approximately 2.3 with 12% PPI scaling is in reasonable agreement with the experimentally observed 3-fold reduction (from 37.1 to 11.9 mPa · s, Table 2). In line with these findings, Tsumura et al. reported for full-length omalizumab that Arg/Cl lowers viscosity by a factor of 2 to 3 compared to Na/Cl, ^55^ consistent with our results for the F(ab’)_2_ fragment. It is encouraging that this trend is captured in the CG-MD simulations.

The GK integrals from the simulations at the F(ab’)_2_ concentrations of 150, 200, and 250 mg/ml with 12% PPI scaling are shown in Figure 4, and the viscosity values are listed in Table 4 (the respective GK curves of the five individual simulations are plotted in Figure S6). For a comprehensive overview, Figure 5 shows the viscosity values from the simulations, the rheology experiments, and their fit to the Ross-Minton equation^56^

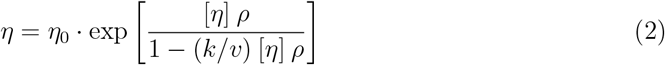

where *η*_0_ is the viscosity of the buffer, [*η*] the intrinsic viscosity of the solute, *ρ* is the mass concentration, *k* is a crowding factor, and *v* is a parameter that depends on the solute shape.

**Table 4:** Dynamic viscosity computed from the CG-MD simulations for different F(ab’)_2_ concentrations and excipient environments.

| $F(ab')_2$ concentration (mg/ml) | Viscosity (mPa · s) | |
| --- | --- | --- |
|  | 150 mM Na/Cl | 150 mM Arg/Cl |
| 150 | $3.4 \pm 0.2$ | $4.8 \pm 0.6$ |
| 200 | $24.4 \pm 6.5$ | $10.5 \pm 1.7$ |
| 250 | $31.7 \pm 14.3$ | $17.2 \pm 2.6$ |

**Figure 4.**
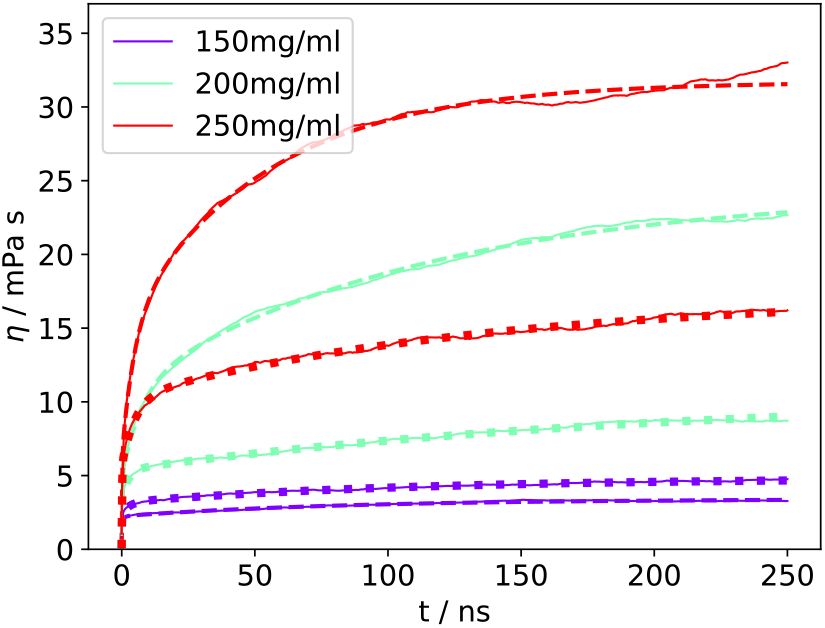
GK integrals from the CG-MD simulations at different F(ab’)_2_ concentrations. The solid lines depict the average GK curves (averaged over five individual simulations), and the dashed and dotted lines show the fits of the GK curves for the systems with Na/Cl and Arg/Cl, respectively

**Figure 5.**
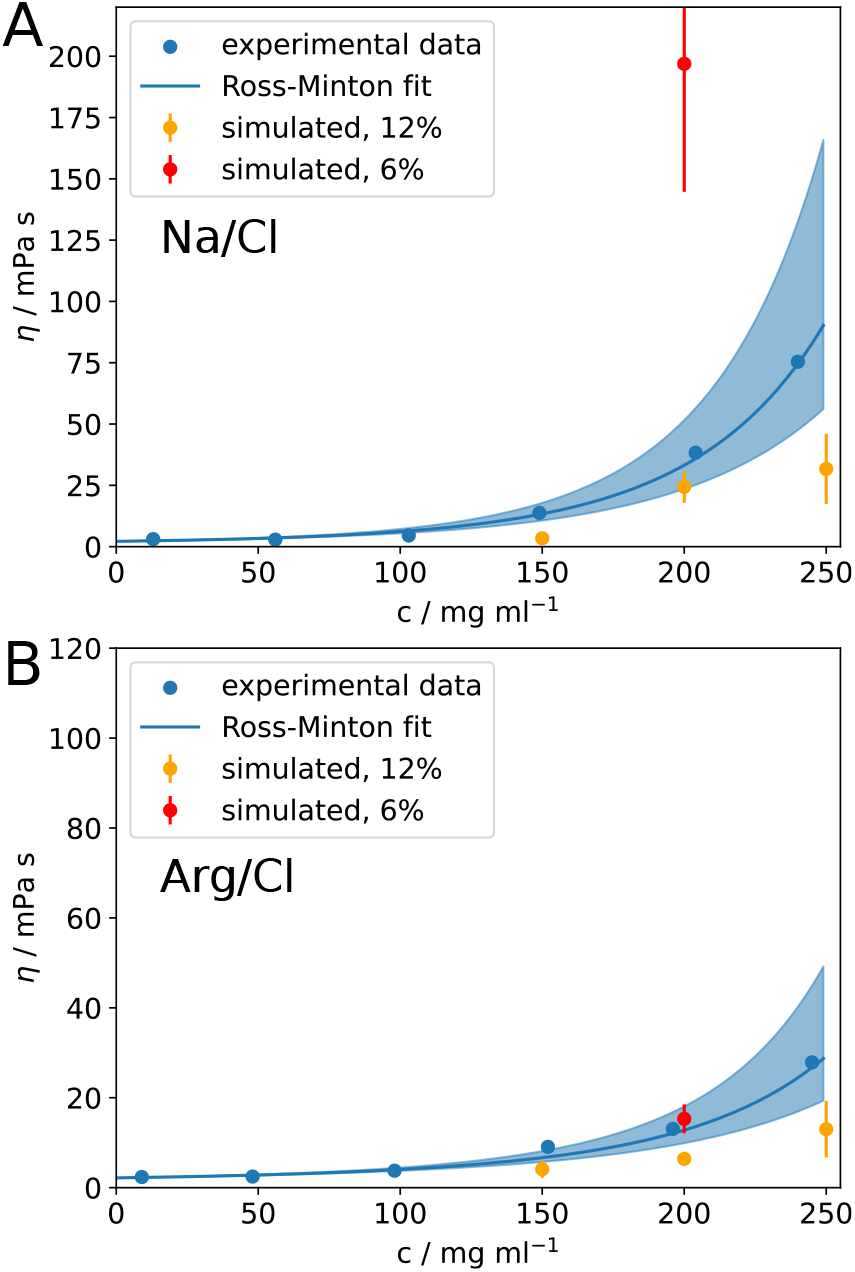
Comparison of dynamic viscosity from rheology experiments and CG-MD simulations. A) and B) show the data for 150 mM Na/Cl and 150 mM Arg/Cl, respectively (note different y-axis scales). Experimental data were fitted with the Ross-Minton equation (Eq. 2). The light blue shaded areas depict an estimated range resulting from an uncertainty in the experimental concentrations of 10%.

Figure 5 shows the expected sharp increase in viscosity with increasing concentration, as predicted by Eq. 2. The viscosities from the CG-MD simulations with 12% PPI scaling are slightly lower than the experimental values, as discussed above. Interestingly, with 6% PPI scaling, the viscosity predicted from the simulation with Arg/Cl lies within the uncertainty of the experimental data, whereas the viscosity with 150 mM Na/Cl is overestimated (red data points in Figure 5). These findings highlight remaining challenges and room for further improvement in the optimization of the Martini 3-exc force field, likely requiring more finetuned adjustments than global scaling of the protein–protein Lennard-Jones interactions.

Finally, we sought to elucidate the molecular basis of the viscosity-lowering effect of arginine excipients. Our previous all-atom MD simulations showed that arginine preferentially associates with charged and aromatic residues, particularly in the CDRs,^33^ consistent with other reports.^31,57^ This suggests that arginine may mask “sticky” surface patches and thereby attenuate attractive PPIs that drive formation of the associative protein network underlying high viscosity at high concentration. To test this hypothesis, we quantified intermolecular contacts between F(ab’)_2_ fragments (F(ab’)_2_–F(ab’)_2_) and contacts involving CDR residues with other F(ab’)_2_ molecules (CDR–F(ab’)_2_). As summarized in Table 5, the total number of F(ab’)_2_–F(ab’)_2_ contacts decreased markedly in the presence of arginine, from 65.0 to 24.9 per F(ab’)_2_ fragment. CDR residues contribute roughly one third of these contacts (19.0/65.0 and 8.8/24.9 for Na/Cl and Arg/Cl, respectively), which is approximately twofold higher than expected purely based on their solvent-accessible surface area (SASA; bottom row of Table 5). Reduced protein–protein contacts are compensated by contacts with arginine: the average numbers of CDR–arginine and F(ab’)_2_–arginine contacts are 16.6 and 41.5 per F(ab’)_2_, respectively (Figure S7). Taken together, these data support a mechanism in which arginine preferentially binds to CDRs and other sticky surface regions, competes with protein–protein association, and thereby loosens the transient protein network responsible for elevated viscosity under the high-concentration conditions studied here.

**Table 5:** Molecular contacts per F(ab’)_2_ molecule between CDR regions and F(ab’)_2_, and between two F(ab’)_2_ fragments in the 200 mg/ml simulations with Na/Cl or Arg/Cl. Molecular contacts were counted between all pairs of CG beads that are within ≤ 0.75 nm from each other (as previously described^52^), and only intermolecular contacts were counted (that is, contacts within a single F(ab’)_2_ were disregarded). The numbers in the table are averages over all molecules in the simulation box and over the final 4 *µ*s of each of the five repeat simulations. The solvent accessible surface areas (SASAs) of the CDRs and the F(ab’)_2_ fragments are 64.0 and 359.0 nm^2^, respectively.

| Molecular contacts | 150 mM Na/Cl |  | 150 mM Arg/Cl |  |
| --- | --- | --- | --- | --- |
| | $F(ab')_2$–$F(ab')_2$ | CDR–$F(ab')_2$ | $F(ab')_2$–$F(ab')_2$ | CDR–$F(ab')_2$ |
| Total contacts | 65.0 | 19.0 | 24.9 | 8.8 |
| Contacts per nm<sup>2</sup> SASA | 0.18 | 0.30 | 0.07 | 0.14 |

Taken together, the results presented in this work underline the suitability of the refined Martini 3-exc force field to study high-concentration antibody solutions and predict excipient effects, at least at a semi-quantitative level. Due to the statistical uncertainty inherent with the viscosity computations, small viscosity differences may not be resolvable by the approach. However, more prominent differences, such as those between Na/Cl and Arg/Cl, can be predicted by the CG-MD simulations, which is promising for formulation design.

In conclusion, this proof-of-principle study demonstrates the feasibility of using coarsegrained MD simulations with the refined Martini 3-exc force field to investigate high-concentration antibody solutions. Our results indicate that the PPI scaling originally introduced by Thomasen et al. for IDPs^40,41^ is also applicable to describe the effective strength of protein– protein interactions in the dynamic protein networks underlying the high viscosity of mAb solutions. The simulations correctly reproduce the trends of increasing viscosity with increasing protein concentration and the viscosity-reducing effect of arginine excipients. However, with 12% PPI scaling, the absolute viscosity values are slightly underestimated. Rather than aiming for fully quantitative agreement of the viscosity values, the CG-MD approach introduced here enables rapid exploration of qualitative trends, and it can provide molecular-level insight into the mechanisms underlying experimental observations. Importantly, the viscosities from the simulations with different excipient environments (150 mM Na/Cl versus 150 mM Arg/Cl) are sensitive to the different nature of the ions, supporting the utility of the CG-MD approach for predicting the effects of different solvation or excipient conditions.

Some limitations remain. In the low-viscosity regime, small differences between different solvation conditions cannot be reliably resolved. In the high-viscosity regime, statistical uncertainties increase and extended simulation times (sampling) are required to observe trends with statistical significance, which can be computationally demanding even at the coarse-grained level.

Future work could extend this approach to a broader range of proteins and excipients to further validate and generalize the findings. CG-MD simulations have the potential to elucidate the molecular mechanisms underlying viscosity modulation by various formulation factors. From a practitioner’s perspective, expanding the library of well-parametrized excipients and additives would facilitate the rapid implementation of the CG-MD methodology, possibly enabling more efficient prescreening of formulation parameters in biopharmaceutical development. Beyond biopharmaceuticals, the CG-MD approach presented here may also have implications for molecular simulations of realistic models of the cell cytoplasm or even whole cells.^58–61^

## Acknowledgement

This work was supported by Boehringer Ingelheim Pharma GmbH & Co. KG and by Deutsche Forschungsgemeinschaft (DFG) under Germany’s Excellence Strategy - EXC 2033 - 390677874 - RESOLV. We thank Kresten Lindorff-Larsen for useful discussions and for hosting T.P. during his internship at the University of Copenhagen.

## Supporting Information Available

GK plots of the 200 mg/ml simulations with Na/Cl at different PPI scalings; GK plots of the 200 mg/ml simulations with all His side chains protonated and GK plots of the GōMartini simulations with 200 mg/ml F(ab’)_2_ concentration; GK plot of the 200 mg/ml simulations with 64 F(ab’)_2_; GK plots of the 200 mg/ml simulations with Arg/Cl; GK plots of the simulations at different mAb concentrations; Molecular contacts of CDR–F(ab’)_2_, F(ab’)_2_–F(ab’)_2_ contacts and CDR–arginine and F(ab’)_2_–arginine contacts; Martini 3-exc force field parameter and coordinate files with starting coordinates of the simulations, MD parameters file, force field topology files, are available at https://doi.org/10.17877/RESOLV-2025-MHT4BTCR

