## Supplementary Information for "Predicting the Dynamic Viscosity of High-Concentration Antibody Solutions with a Chemically Specific Coarse-Grained Model"

### Methods

#### MD Simulations

The atomic coordinates of the omalizumab Fab domain were obtained from the protein data bank (PDB: 2XA8).[S1] The structural model of the omalizumab F(ab')$_2$ was generated using MODELLER,[S2,S3] with the equilibrated trastuzumab structure from Brandt et al.[S4] serving as a template. The 2XA8 structure was superimposed onto both Fab parts of the template, and the heavy chain was extended using the template structure (hinge and Fc domain). Afterwards, the full-length mAb was cleaved after H-C232, i.e., after the second disulfide bridge in the hinge region. The Martinize2 script[S5] was used to convert the atomic coordinates to a coarse-grained Martini 3 representation and to generate a force field topology, including the disulfide bridges. The side chain corrections of Herzog et al.[S6] were applied. Furthermore, the OLIVES[S7] model was applied to the F(ab')$_2$ structures, with the recommended settings. OLIVES potentials between the two separate Fab domains of the F(ab')$_2$ dimer were removed except for the ones in the hinge region. In addition, we carried out control simulations with the recent GōMartini model[S8] instead of OLIVES. The GōMartini restraints were generated with Martinize2,[S5] using the default settings as described by Souza et al.[S8]

For the CG-MD simulations, cubic simulation boxes containing 32 F(ab')$_2$ molecules were set up with protein concentrations of 150, 200, and 250 mg/ml. 150 mM Na/Cl or 150 mM Arg/Cl were added to each simulation box (2904, 2178, or 1742 of each ion species, respectively). The total number of CG beads in the 150, 200, and 250 mg/ml systems were approximately 320.000, 250.000, and 210.000, respectively. Additional control simulations at 200 mg/ml were performed with a twofold larger system containing 64 F(ab')$_2$ molecules (approximately 500.000 CG particles). For every combination (mAb and ion concentration), five independent starting configurations were generated by different random initial placement of the F(ab')$_2$ molecules in the simulation boxes (but no clashes).

MD simulations were performed with the GROMACS simulation package (version 2021.1)[S9] using the Martini 3 force field.[S10] To adjust the PPIs, all interactions between protein beads were scaled by reducing the well-depth $\epsilon$ of the Lennard-Jones (LJ) 6,12 potential (that is, making the LJ interactions less attractive) by 6%, 12%, 24%, or 36%. The simulation systems were energy minimized and afterwards equilibrated for 50 ns in the isobaric-isothermal ensemble with position restraints on the proteins at a constant temperature of 300 K and constant pressure of 1 bar, using the velocity rescaling thermostat with a stochastic term[S11] and the stochastic cell rescaling barostat,[S12] respectively. The recommended "new-rf"[S13] simulation settings were used with 20 fs time steps and reaction field electrostatics. Additionally, the recommended neighbor list settings by Kim et al. were applied.[S14]

The final production simulations were performed in the canonical (NVT) ensemble for 5 $\mu$s each, yielding a total sampling time of 25 $\mu$s for every system. The starting configurations of the production runs were chosen by computing the average simulation box volume of the equilibration simulations, taking the last equilibration frame with a box volume smaller than the average, and setting the box volume of that frame to the average.

#### Viscosity Computation

For computing the dynamic viscosity from the MD simulations, we followed the approach described in detail in our previous work[S15] and computed the dynamic viscosity from the Green-Kubo (GK) integrals (equation 1 in the main text). In addition to the nondiagonal elements $P_{xy}$, $P_{yz}$, and $P_{xz}$, we used the following combinations of diagonal elements, $(P_{xx} - P_{yy})/2$, $(P_{xx} - P_{zz})/2$, and $(P_{yy} - P_{zz})/2$, to improve statistics.[S16,S17] To capture the fast pressure fluctuations, the pressure tensor was saved to disk every 20 fs. The pressure-pressure ACFs were computed up to a maximum lag time of 500 ns (10% of the trajectory length) and averaged over all five independent 5 $\mu$s simulations. The resulting GK integrals were fitted with a triexponential function

$$\eta(t) = B \cdot \alpha \cdot \tau_1 \left(1 - e^{-t/\tau_1}\right) + B \cdot \beta \cdot \tau_2 \left(1 - e^{-t/\tau_2}\right) + B \cdot (1 - \alpha - \beta) \cdot \tau_3 \left(1 - e^{-t/\tau_3}\right) \quad (1)$$

with fitting parameters $B > 0$, $\alpha, \beta < 1$, and $\tau_{1,2,3} > 0$. The GK integrals were fitted with equation 1 up to a maximum lag time of 250 ns. As previously described,[S15] the data were non-uniformly weighted in the fit (according to the standard deviation), so that the statistically more precise data at shorter lag times contribute stronger than noisy data at long lag times. The viscosity was determined from the asymptotic limit ($t \to \infty$) of the analytical fit function. The code used for these calculations is available on GitHub (`https://github.com/MolSimGroup/gmx_gk_autocorr/`).

#### mAb and F(ab')$_2$ Production, Purification and Formulation

The omalizumab monoclonal antibody (mAb) used in this study was supplied by Boehringer Ingelheim Pharma GmbH & Co. KG (Biberach an der Riß, Germany). Omalizumab, an IgG1 isotype with an approximate molecular weight of 150 kDa, was produced via mammalian cell culture in Chinese Hamster Ovary (CHO) suspension-adapted cells and purified as described previously.[S15] Purification involved affinity, ion-exchange, and size-exclusion chromatography, with final formulation into 10 mM histidine buffer pH 6.5 via preparative size-exclusion chromatography.

For F(ab')$_2$ fragment generation, omalizumab was enzymatically cleaved using FabRICATOR™ (Genovis, Kävlinge, Sweden) following the manufacturer's protocol with modifications for large-scale processing. The digested sample was diluted in PBS and subjected to preparative Protein A affinity chromatography (Thermo Scientific), isolating F(ab')$_2$ in the flow-through and neutralizing with Tris buffer pH 8. The F(ab')$_2$ fragment was further polished and formulated in 10 mM histidine buffer pH 6.5 using size-exclusion chromatography. Purity and identity of omalizumab mAb and F(ab')$_2$ were confirmed by size-exclusion chromatography, non-reduced capillary gel electrophoresis, and liquid chromatography-mass spectrometry (LC-MS).

F(ab')$_2$ were concentrated to 100 mg/mL in 10 mM histidine buffer pH 6.5. Aliquots were spiked with sodium chloride (Na/Cl) or arginine hydrochloride (Arg/HCl) to prepare the following formulations: (i) 10 mM histidine pH 6.5 (non-spiked), (ii) 10 mM histidine + 150 mM Na/Cl pH 6.5, and (iii) 10 mM histidine + 150 mM Arg/HCl pH 6.5 with pH adjustments with concentrated base or acid as required. Samples were concentrated to 250 mg/mL or the maximum achievable concentration and subsequently diluted to 200, 150, 100, 50, and 10 mg/mL in the respective formulation buffers (i, or ii, or iii). Protein concentrations were verified using variable pathlength slope spectroscopy (SoloVPE; C Technologies Inc., New Jersey, USA), and the measured concentrations were used for rheological characterization.

#### Rheology Experiments

Dynamic viscosity measurements were conducted at 20 °C using a RheoSense VROC Initium One Plus rheometer (San Ramon, CA, USA) equipped with a B05 chip. A 40 $\mu$l sample volume was loaded into deactivated clear glass HPLC vials (Waters). The viscosity was evaluated across a shear rate range of 250 $s^{-1}$ to 2000 $s^{-1}$, with measurements recorded at 250 $s^{-1}$, 500 $s^{-1}$, 1000 $s^{-1}$, 1500 $s^{-1}$, and 2000 $s^{-1}$. Data were analyzed for non-Newtonian behavior (e.g., shear thinning) or potential interfacial interaction artifacts. Dynamic viscosity was measured at a fixed shear rate of 1000 $s^{-1}$ in six replicates per sample at 20 °C, and the mean values were reported. System suitability testing (SST) was performed using a medical-grade viscosity standard from Paragon Scientific (ISO 17025 and 17034 certified), with a reported dynamic viscosity of 9.994 mPa·s and a density of 1.1567 g/ml at 25 °C.

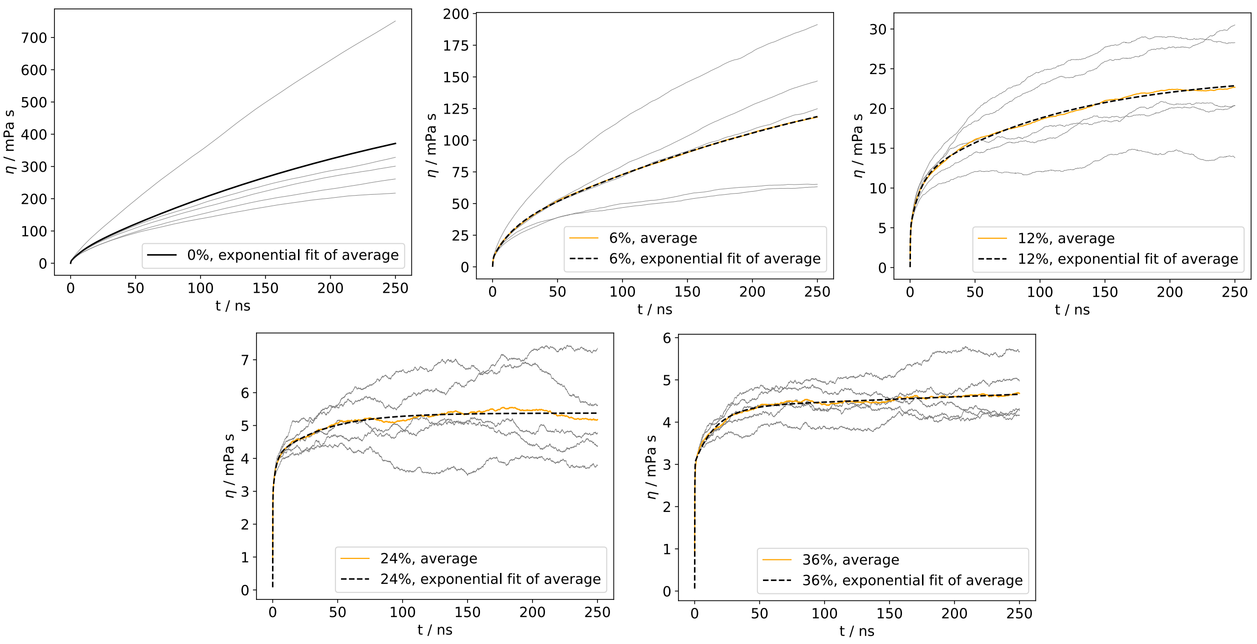


Figure S1: Green-Kubo integrals from the 200 mg/ml omalizumab $F(ab')_2$ simulations with 150 mM Na/Cl and different PPI scalings. The triexponential fits to the average GK integrals are shown as dashed lines.

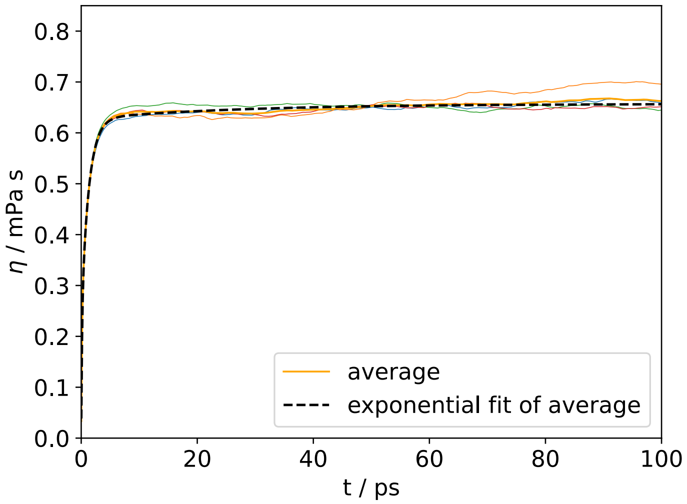


Figure S2: Green-Kubo integrals from 40 ns simulations of a box with 10000 water Martini 3 water beads at 300 K. The dynamic viscosity obtained from the asymptotic limit is 0.66 $\mathrm{mPa} \cdot \mathrm{s}$.

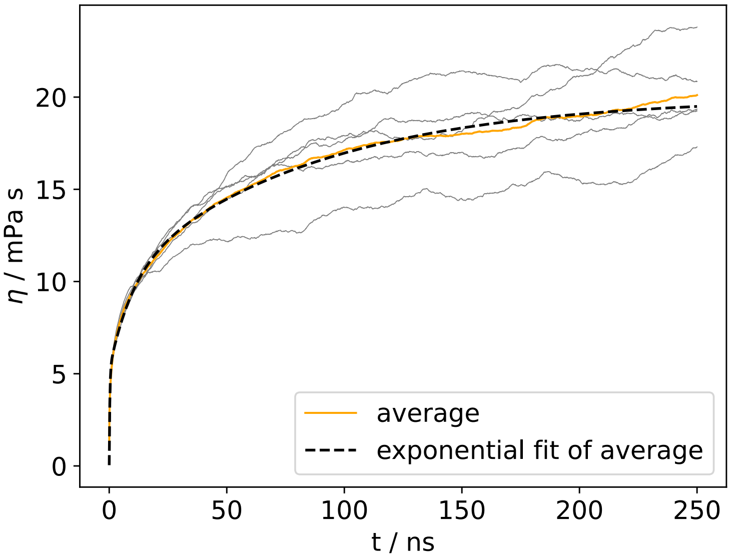


Figure S3: Green-Kubo integrals from simulations with 64 omalizumab $\mathrm{F(ab')_2}$ domains at 200 mg/ml with 150 mM Na/Cl. The dynamic viscosity obtained from the asymptotic limit is $20.0 \pm 2.3$ $\mathrm{mPa} \cdot \mathrm{s}$.

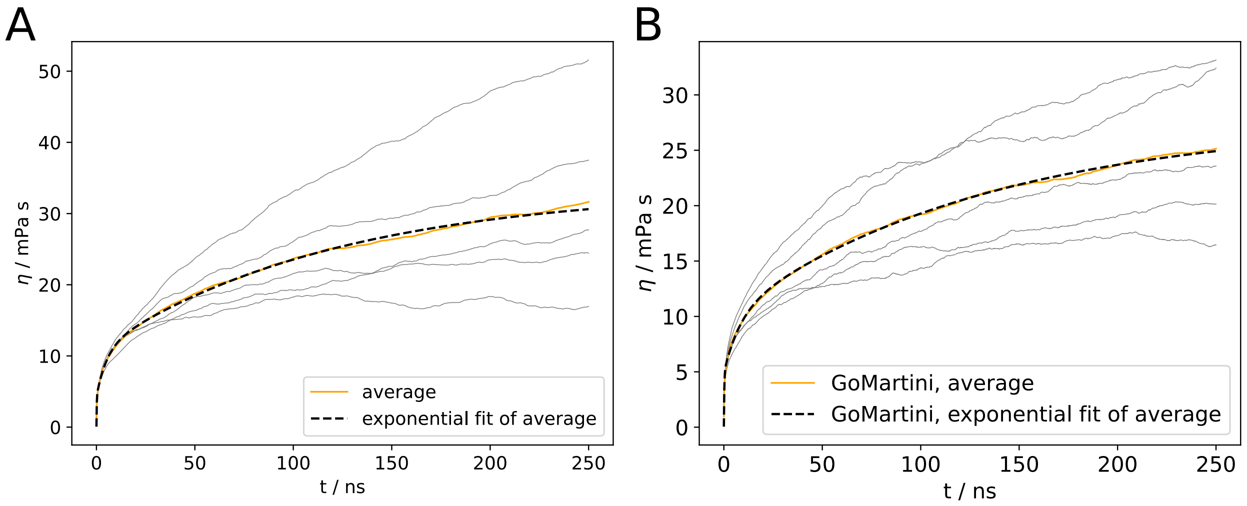


Figure S4: Green-Kubo integrals from 200 mg/ml omalizumab F(ab')$_2$ simulations with A) 150 mM Na/Cl, in which all His side chains were modeled as positively charged (that is, doubly protonated), and B) 150 mM Na/Cl with the GōMartini 3 model[S8] instead of OLIVES. The asymptotic limits of the GK integrals yield viscosities of 33.5 ± 17.3 and 27.6 ± 8.5 mPa · s, respectively.

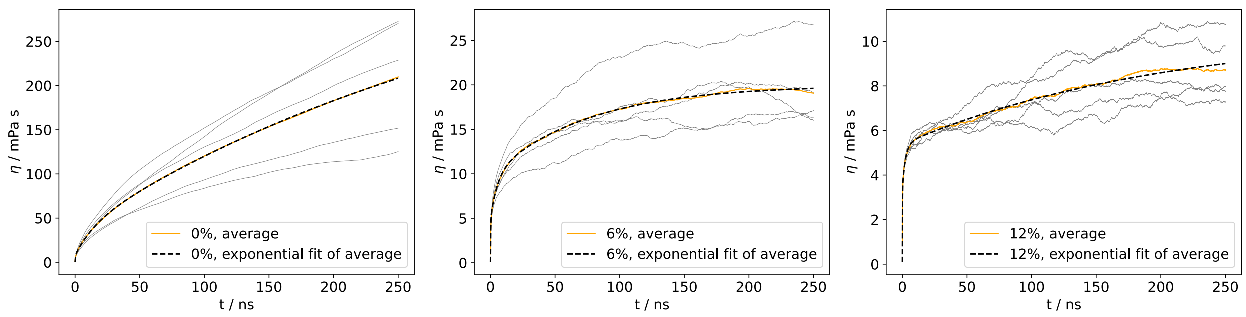


Figure S5: Green-Kubo integrals from the 200 mg/ml omalizumab F(ab')$_2$ simulations with 150 mM Arg/Cl and PPI scalings of 0% (left), 6% (middle), and 12% (right).

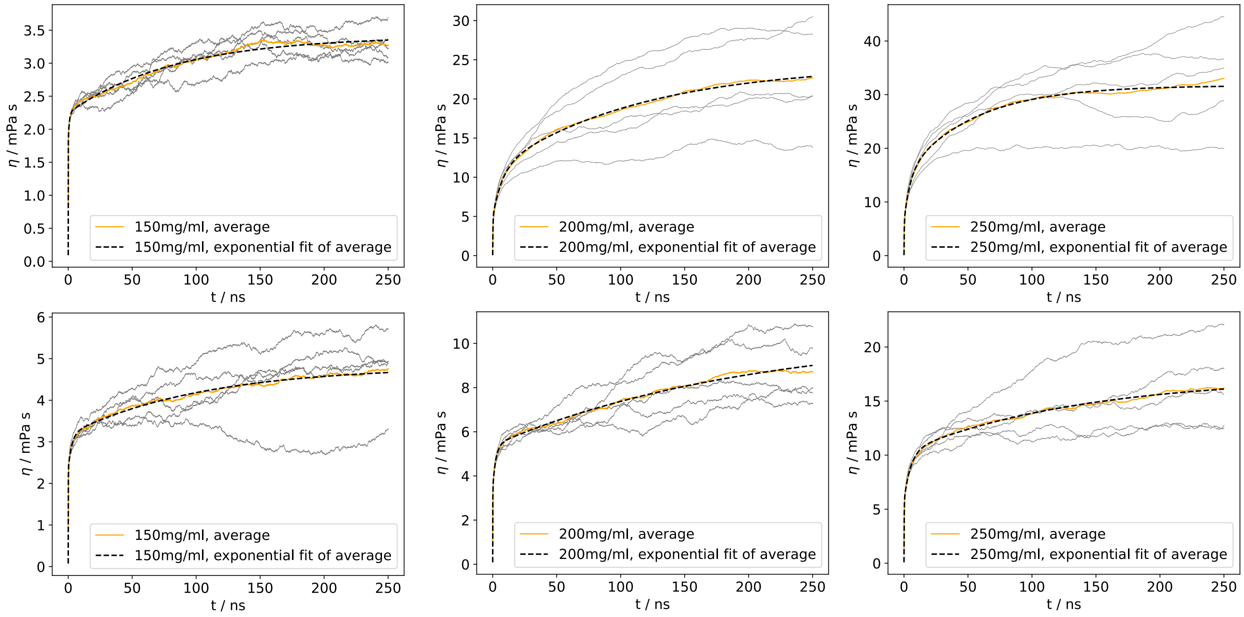


Figure S6: Green-Kubo integrals from the omalizumab F(ab')$_2$ simulations at concentrations of 150, 200, and 250 mg/ml with 150 mM Na/Cl (top row) and 150 mM Arg/Cl (bottom row) at PPI scaling of 12%.

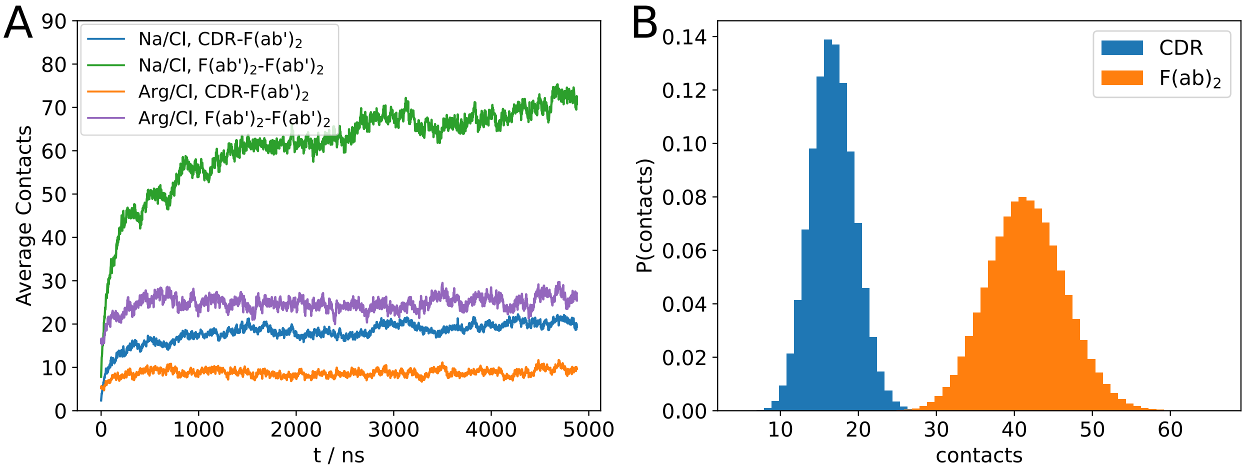


Figure S7: Molecular contacts (counted within 0.75 nm bead–bead distance) in the 200 mg/ml F(ab')$_2$ simulations with Na/Cl and Arg/Cl. A) shows the system average (over time and all F(ab')$_2$) CDR–F(ab')$_2$ and F(ab')$_2$–F(ab')$_2$ molecular contacts, where only intermolecular contacts are considered. Counted are all CG beads in molecular contact distance of the reference CDR or F(ab')$_2$. B) shows the contact histograms of the CDR or F(ab')$_2$ domains with arginine excipients in the 200 mg/ml F(ab')$_2$ simulations with Arg/Cl, where the number of arginine excipient molecules were counted. One contact with arginine was counted if any CG bead of an arginine excipient molecule was within 0.75 nm of any bead of the CDRs (blue histogram) or any bead of the F(ab')$_2$ (orange histogram). Thus, the CDR data is a subset of the F(ab')$_2$ data and includes all CDR loops in both heavy and light chains of the two individual Fab domains of each F(ab')$_2$ dimer.
